# Peptide frontier-orbital gaps narrow by backbone electrostatics

**DOI:** 10.64898/2026.10.05.756722

**Authors:** Xu Wu, Wei Fu

## Abstract

Aromatic side chains carry the optical and redox chemistry of peptides, and lengthening one is widely expected to extend its conjugation and narrow its HOMO-LUMO gap. We test that expectation with plane-wave density functional theory on 22 isolated neutral peptides NH_2_-(X)_n_-COOH (X = Phe, Tyr, Trp, Cys; n = 1 to 5, and n = 10 for Cys and Trp). The gap does narrow, at the same rate for all three aromatic residues, i.e., −0.17 to −0.18 eV per residue, even though benzene, phenol and indole are very different chromophores. Residue-resolved projections reveal why: nothing delocalises. Every aromatic frontier orbital is 97 to 100 % confined to a single residue and the HOMO participation number saturates at one ring. Each residue instead keeps its own local gap: for Trp it is invariant to within 0.03 eV over n = 2 to 10, while over that same span the global gap falls by 1.19 eV. The global gap is therefore an extreme-value functional of the residue site energies, gap(n) = ⟨*g*_*f*_⟩ − Δ(n), whose alignment penalty accounts for the entire effect: the polar backbone generates a monotonic internal electrostatic ramp that leaves the two gap edges at opposite ends of the chain, 32 Å apart. We set out the resulting design rules for peptide-based bio-organic semiconductors.

## 1. Introduction

Peptides and proteins are increasingly proposed as the active component of bioelectronic devices, e.g., as charge-transporting layers, as the recognition element in biosensors, and as the self-assembling scaffold of conductive fibrils and nanotubes [1–4]. For any such material the quantity that governs injection and transport is the alignment of its frontier-orbital levels, and the parameter that is easiest to vary is chain length. The intuitive expectation is that a longer peptide means a longer conjugated path, a smaller HOMO–LUMO gap and a redshifted absorption edge. That expectation is repeated often enough to have become the default, and it is the one against which the present results were initially read.

It deserves scrutiny, because the peptide backbone is a poor conductor of π conjugation. Each residue contributes an amide unit whose nitrogen lone pair is conjugated with the carbonyl, but successive amide π systems are separated by an sp^3^ α-carbon bearing two substituents, and the amide-to-amide dihedrals φ and ψ are variable. This is why the peptide bond is described as a *localised* amide resonance rather than a conjugated polymer repeat unit [5], and why the classic picture of a peptide is that of a spacer, i.e., an insulating bridge whose length tunes electron transfer between remote donors and acceptors [6, 7].

If the backbone neither extends conjugation nor delocalises frontier orbitals, what does chain length actually do? We answer that by direct computation, and find the answer is electrostatic rather than electronic in the usual sense. The systems are small enough (up to 243 atoms) for well-converged plane-wave DFT, they are isolated so the comparison is free of solvation and packing effects, and the LORBIT = 11 atom projection lets us ask not merely *what* the gap is but *where each frontier orbital lives*. That combination, a residue-resolved decomposition compared explicitly against the intra-residue gap, is what separates the two candidate mechanisms, and it is the main methodological contribution of this work.

We include cysteine as a deliberate control. Cysteine is non-aromatic: sulfur lone pairs and the backbone dominate its frontier orbitals. If the aromatic trend is a chromophore effect, cysteine should not share it; if it is a backbone effect, cysteine should, albeit with the caveat that its frontier orbitals differ in character, which leads instead to genuinely different behaviour.

## 2. Results and discussion

### 2.1 The gap narrows at a rate that does not depend on the chromophore

For the three aromatic residues the gap falls monotonically and approximately linearly with n (Figure 1):

**Figure 1.**
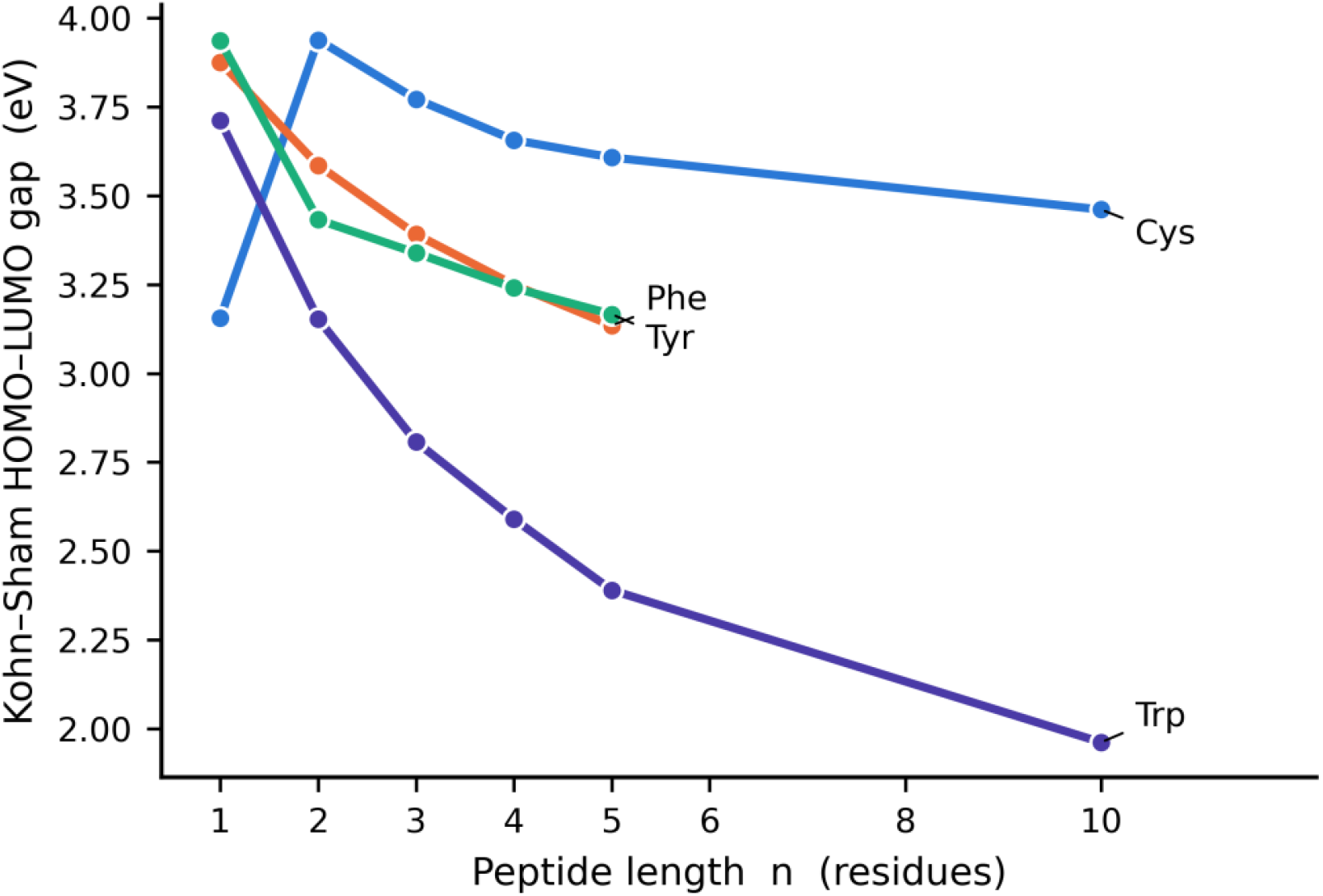
Kohn–Sham HOMO–LUMO gap versus peptide length n for the four residues. Phe, Tyr and Trp fall along nearly parallel lines; cysteine is flat.

| Residue | Range | Slope (eV per residue) | R |
| --- | --- | --- | --- |
| Phe | $n = 1$ to 5, 3.876 → 3.135 eV | $-0.182 \pm 0.020$ | -0.982 |
| Tyr | $n = 1$ to 5, 3.936 → 3.167 eV | $-0.173 \pm 0.048$ | -0.901 |
| Trp | $n = 1$ to 10, 3.712 → 1.962 eV | $-0.174 \pm 0.040$ | -0.910 |
| Cys | $n = 1$ to 10, 3.156 → 3.461 eV | $-0.007 \pm 0.042$ | -0.079 |

Uncertainties are standard errors of the least-squares slope. The scatter about the fitted lines is not negligible (residual standard error 0.28 eV for Trp), so the robust comparison is between slopes rather than against zero. The three aromatic slopes agree well within their uncertainties (Phe and Trp differ by 0.008 eV per residue against a combined standard error of 0.045), whereas cysteine is smaller than each by a factor of about 26.

This is the result that redirects the analysis. **The three aromatic residues narrow their gaps at the same rate**, yet benzene, phenol and indole have markedly different π systems (six, six-plus-oxygen and nine ring atoms) and quite different gaps: Trp contains an indole, a substantially better electron donor than benzene, and one-residue Trp has a gap of 3.71 eV against 3.88 eV for Phe. If the trend came from π systems coupling along the backbone, the slope would scale with the chromophore’s π extent and with the inter-residue coupling. It does not. A trend insensitive to the identity of the chromophore is unlikely to be a property *of* the chromophore.

Both gap edges contribute to the change rather than one carrying it: for Trp the HOMO rises by 1.31 eV while the LUMO falls by 0.44 eV, and for Phe the two are of similar size and opposite sign (Figure S2), so a mechanism that shifts the frontier manifold rigidly cannot be the whole story.

### 2.2 Nothing delocalises

The most direct test of a conjugation mechanism is whether the frontier orbitals grow (Figure 3). Summing the per-atom weights over residues, every aromatic system with n ≥ 2 carries **at most 2.0 %** of its HOMO weight outside the hosting residue and **at most 2.5 %** of its LUMO weight (Table S4). The host residues carry 97.5 to 100 % of the density (Table 1).

**Table 1.** Where the global HOMO and global LUMO actually live.

| System | HOMO host residue | weight | LUMO host residue | weight | separation (Å) |
| --- | --- | --- | --- | --- | --- |
| $(W)_2$ | 2 | 0.99 | 1 | 1.00 | 8.5 |
| $(W)_3$ | 2 | 0.99 | 3 | 1.00 | 7.6 |
| $(W)_4$ | 2 | 0.99 | 3 | 1.00 | 13.2 |
| $(W)_5$ | 2 | 0.99 | 5 | 1.00 | 14.5 |
| $(W)_{10}$ | 2 | 0.98 | 9 | 1.00 | 32.7 |
| $(F)_5$ | 1 | 0.98 | 5 | 0.98 | 16.4 |
| $(Y)_5$ | 1 | 0.99 | 5 | 0.99 | 15.6 |
| $(C)_{10}$ | 1 | 0.97 | 10 | 0.62 | 31.7 |

The participation number tells the same story (Figure 4). For Trp it rises only from 7.95 at n = 1 to 8.95 at n = 10, saturating at the ∼9 atoms of a single indole ring and going no further. For Phe it stays between 1.8 and 2.6 atoms (1.76 to 2.52), and for Tyr between 4.2 and 4.8, with a single outlier at 6.7 for (Y)_2_; there the same orbital is spread more evenly over its ring rather than concentrated on fewer atoms, and it remains 99.3 % on the residue (Table S4), so the outlier is a change of shape and not of locality. The orbitals are not becoming more extended; if anything they become slightly more compact as the surrounding polar backbone polarises them.

Two conclusions follow, neither of them what the initial intuition suggested. There is **no extended conjugation** in these peptides at any length studied; and chain length is not changing what the frontier orbitals *are*, but something about where they *sit*.

### 2.3 The decisive measurement: the local gap does not move

If the chromophores are untouched, the gap of a single residue measured *within* that residue must be independent of n, even while the global gap collapses. We therefore computed each residue’s own frontier pair (*H*_*f*_, *L*_*f*_) and its local gap *g*_*f*_ = *L*_*f*_ − *H*_*f*_, and compared the mean ⟨*g*_*f*_⟩ against the global gap.

For tryptophan, the longest chain and hence the strongest test, the mean local gap is 3.671, 3.642, 3.648, 3.641 and 3.643 eV at n = 2, 3, 4, 5 and 10. Over the same span, in which the global gap falls by 1.19 eV (3.153 → 1.962 eV), the local gap moves by 0.028 eV, i.e., a drift of 2.4 %. The local gap is, for practical purposes, a constant of the system.

For Phe and Tyr it does not merely stay flat but *widens* slightly (+0.20 and +0.08 eV), the opposite sign to the global trend. This is physically reasonable: a longer peptide is a larger, more polarisable object, and its residues sit in a more screened environment, which slightly raises the penalty for local charge separation within a residue.

#### None of the observed chain-length dependence is a change in any residue’s own excitation energy

The whole trend is a change in the relative alignment of residues’ levels, i.e., an inter-site, charge-transfer-like gap rather than a local one.

This conclusion is insensitive to the threshold that defines “localised on a residue”. Because the aromatic frontier orbitals are 97 to 100 % localised, the assignment of *H*_*f*_ and *L*_*f*_ barely depends on it. Scanning the threshold from 0.15 to 0.60 (Tables S2a and S2b), the ⟨*g*_*f*_⟩ drift for Trp between n = 2 and n = 10 stays between −0.028 and +0.154 eV against a global drift of −1.192 eV, at most 13 % of the effect, and only +2 % at the working threshold of 0.45. Tyr reaches 29 % and Phe 44 % at that threshold, and more at coarser ones; Phe is the extreme case because its local gap genuinely *widens* with n (+0.197 eV, Table 2), the opposite sign to the global trend. In no case does the local drift carry the global trend, and for the decisive long-chain case, Trp, it is negligible. At the working threshold every residue of every aromatic system with n ≥ 2 hosts its own well-defined frontier pair, so no residue is being discarded.

**Table 2.** Mean intra-residue gap versus the global gap. Local-gap drifts are measured from n = 2 (at n = 1 the local and global gaps coincide); global-gap drifts are from n = 1.

| Residue | $\langle g_f \rangle$ at $n = 2$ | at $n = 10$ | drift | global gap drift |
| --- | --- | --- | --- | --- |
| Trp | 3.671 eV | 3.643 eV | −0.028 eV | −1.750 eV |
| Tyr | 3.980 ($n=2$) | 4.057 ($n=5$) | +0.077 eV | −0.770 eV |
| Phe | 4.353 ($n=2$) | 4.551 ($n=5$) | +0.197 eV | −0.741 eV |

### 2.4 The global gap is an extreme-value functional of the site energies

If each residue carries its own frontier pair, the global Kohn–Sham gap is

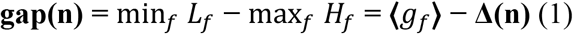

where Δ(n) is an alignment penalty that vanishes only if a single residue hosts both global edges. This is an *extreme-value* (order-statistics) functional: the global HOMO is the highest of the n site HOMOs and the global LUMO the lowest of the n site LUMOs. Equation (1) is an identity and so explains nothing by itself, but it localises the question: we can measure Δ(n) and ask which term carries the chain-length dependence (Table S5).

The answer is total. For Trp between n = 2 and n = 10 (Table 3), ⟨*g*_*f*_⟩ changes by −0.03 eV while Δ(n) grows by +1.16 eV and the gap falls by 1.19 eV. **Δ(n) accounts for the entire effect, indeed slightly more** than the entire effect, since the small increase in the local gap works against the trend. By n = 10, Δ is 46 % of the local gap: the global gap of the decamer, 1.962 eV, is only 56 % of the *smallest* of the ten local gaps it contains (3.517 eV).

**Table 3.**
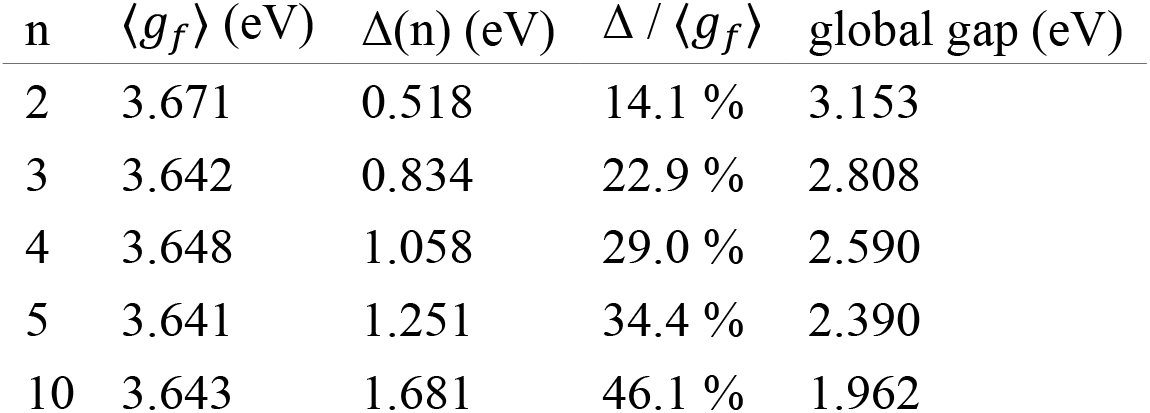
Decomposition of the Trp gap by equation (1).

This also resolves the puzzle of Section 2 Since ⟨*g*_*f*_⟩ is a property of the chromophore but Δ(n) is not, and ⟨*g*_*f*_⟩ carries no n-dependence at all, the entire measured slope must come from Δ(n), which is built from the *backbone* environment and is therefore nearly the same for benzene, phenol and indole. The chromophore-independent slope is a direct consequence of the trend not being a chromophore property.

### 2.5 The origin of Δ: a monotonic electrostatic ramp along the backbone

Why do the site energies spread out as n grows? The spread could be a systematic ramp, i.e., a monotonic shift with position requiring a directed field along the backbone, or statistical disorder, random variation of each site’s environment, in which case Δ(n) would be a pure order-statistics effect growing with n simply because more samples reach further into the tails of a fixed distribution. The two have different physical content, so we distinguish them by regressing each residue’s own frontier energies against its position.

The answer is unambiguous: it is a **ramp** (Figure 6). For (W)_10_ the site HOMO correlates with residue index at R = −0.903 and the site LUMO at R = −0.902; for (W)_5_, R = −0.893 and −0.924; for (Phe)_3_, R = −0.999. These are fits of orbital energy against residue index and are distinct from the gap-versus-separation correlations quoted at the end of this section; the Trp pair coincides numerically at −0.903 only by coincidence. Fitted slopes reach −0.61 eV per residue.

The origin of a monotonic potential along a chain of identical residues is the **permanent electrostatic field of the peptide itself**. The backbone is strongly polar: each amide carries a substantial dipole, the carbonyl oxygens and amide hydrogens are well separated in charge, and the two termini are themselves polar. Because the backbone geometry imposes a preferred directionality on the amide dipoles, these contributions do not cancel but superpose into a net internal potential varying monotonically along the molecular axis. Each pendant chromophore sits at a different point on that potential and its orbital energies are rigidly shifted accordingly. In the language of molecular electronics, the peptide backbone acts as a **molecular electrostatic gate**, and chain length controls how far apart in gate voltage two residues can be (Figure 7).

The fitted per-residue slope is not constant in n; it dilutes as the chain lengthens: for Trp from −0.41 eV per residue at n = 3 to −0.16 at n = 10. But the *total* potential drop across the molecule, the fitted slope multiplied by the number of residue spacings, is roughly constant: 0.94 to 1.41 eV for Phe, 0.40 to 0.80 eV for Tyr, 0.64 to 1.43 eV for Trp, with no systematic trend in n. The potential difference across the molecule is thus approximately a fixed property of its polar groups, and a longer chain simply distributes the same total drop over more residues, i.e., the per-residue field dilutes as roughly 1/n.

This is consistent with the molecules folding as they grow (Figure 2), so that end-to-end distance does not scale linearly with residue count, as is evident from d(H–L) reaching only 32 Å at n = 10 (Table S6). It also means the extreme-value argument does not require the per-residue field to persist at constant strength: a fixed total drop spread over a growing number of sites leaves the *average* site-energy spacing roughly constant while Δ(n) still grows, because the outermost sites reach further into the tails.

**Figure 2.**
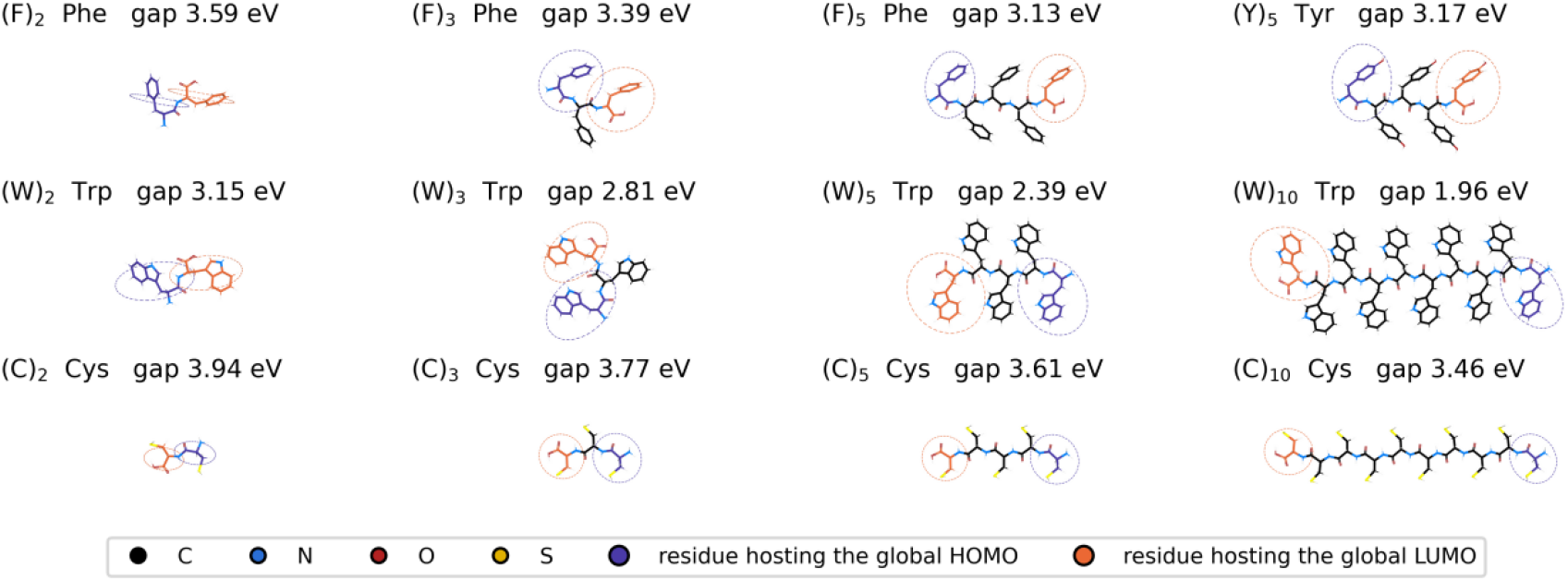
The relaxed geometries, ray-traced in PyMOL with sticks coloured by element (C black, N blue, O red, S yellow), each panel labelled with its Kohn–Sham gap. The residue carrying the global HOMO is picked out in violet and the one carrying the global LUMO in orange, each enclosed in a dashed ring of the same colour. Every panel shares one Å-per-pixel scale, so chain growth is shown at true relative size. From n = 2 onward the two rings never enclose the same residue. For Cys the rings mark the largest weight only: as Section 2.6 shows, the cysteine LUMO is genuinely spread over the backbone and no single residue owns it.

**Figure 3.**
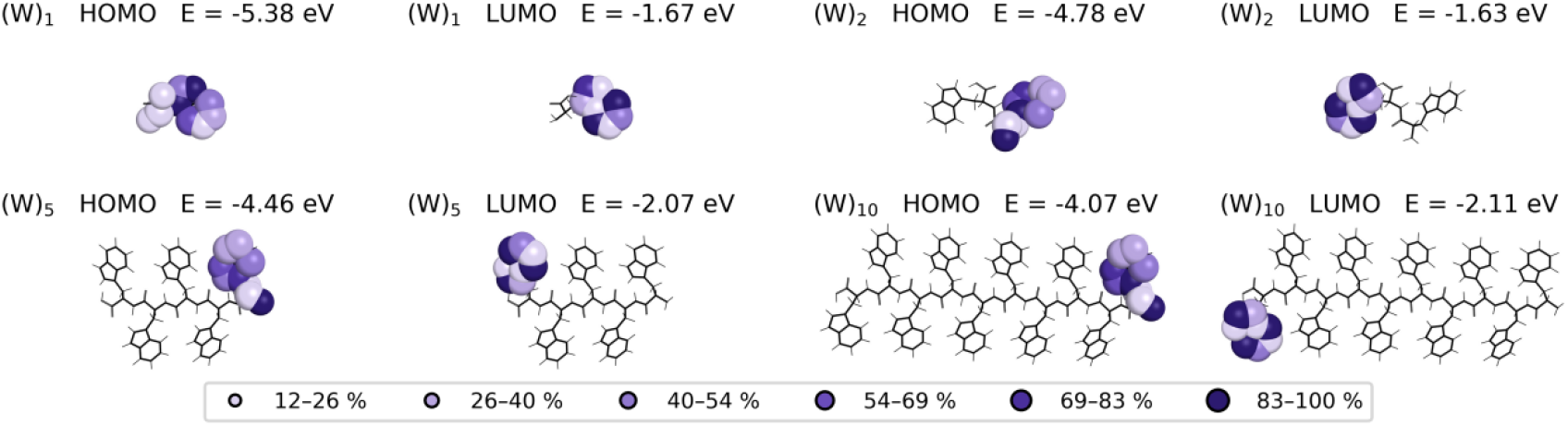
Frontier-orbital densities for (W)_1_, (W)_2_, (W)_5_ and (W)_10_, ray-traced in PyMOL. The skeleton is drawn as black sticks and the orbital as spheres whose radius and shade scale with the per-atom weight from the LORBIT = 11 projection (pale ≈ 12 %, dark = 100 %). All panels share one Å-per-pixel scale. Each orbital is a compact single-ring object that does not spread as n grows.

**Figure 4.**
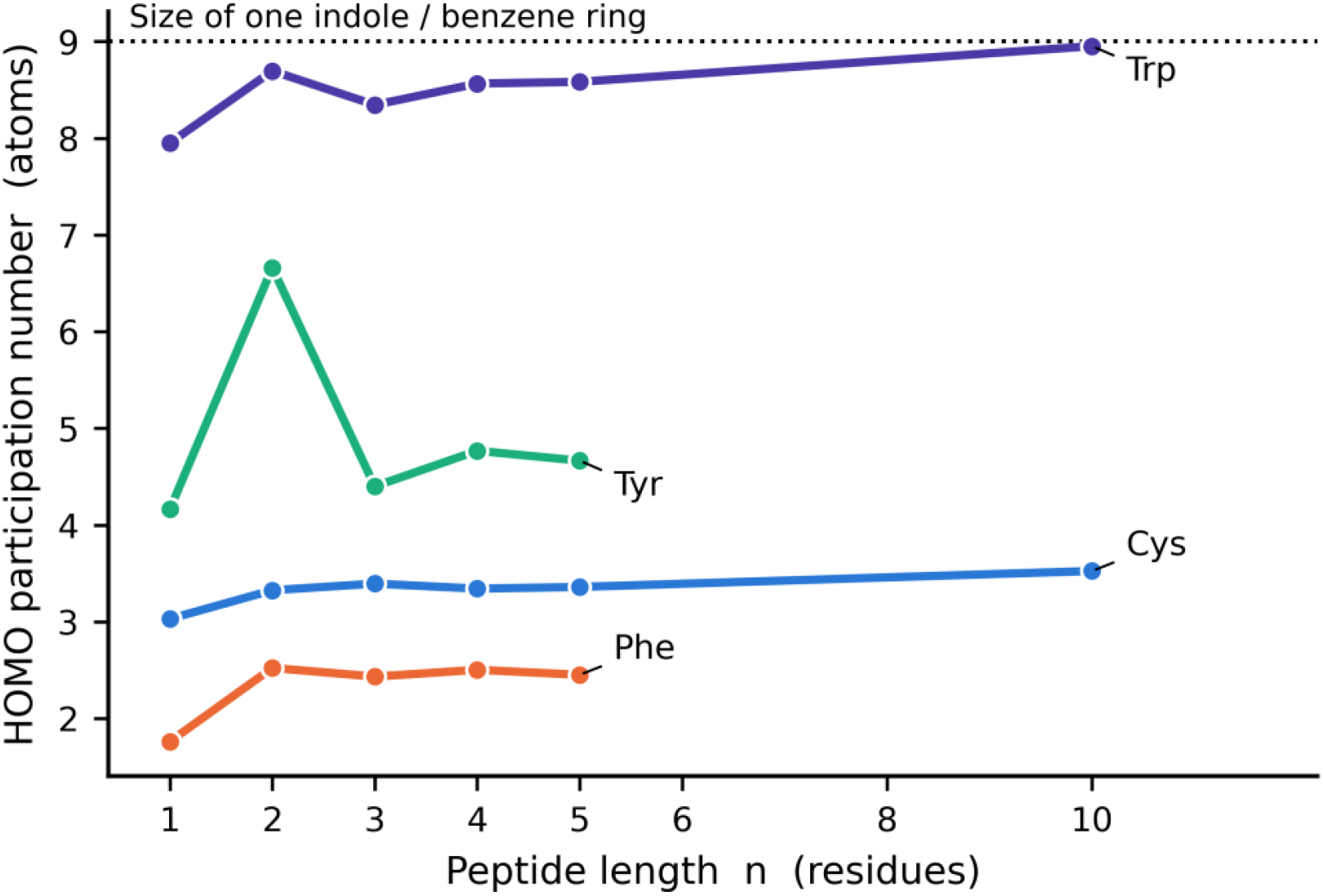
HOMO participation number versus n. The Trp curve saturates at ∼9 atoms, the size of one indole ring (dashed line). No curve shows the growth delocalisation would require.

The correlation between the gap and the physical separation of the edges is strong (Figure S1): gap versus d(H–L) gives R = −0.967 (Phe), −0.905 (Tyr) and −0.903 (Trp) across all n, and −0.988, −0.984 and −0.896 when the n = 1 points are excluded. The gap falls as the two edges are pulled apart, the signature of a spatially indirect gap.

### 2.6 The non-aromatic control: cysteine

Over n = 1 to 10 the cysteine gap is essentially flat, with a fitted slope of −0.007 ± 0.042 eV per residue (R = −0.079), i.e., indistinguishable from zero and about a factor of 26 smaller than the aromatic slope. The control evidence is compositional rather than statistical, and has two parts.

First, the cysteine HOMO is not a ring π state at all: it is dominated by sulfur and nitrogen lone pairs, and changes character across the series (a thiol lone pair at −4.571 eV for n = 1, a backbone nitrogen lone pair at about −5.3 eV for n ≥ 2). Second, and decisively, the cysteine LUMO is substantially **delocalised over the backbone**, whereas no aromatic LUMO is: the off-residue LUMO weight is 0.160 at n = 2, rising monotonically to 0.377 at n = 10, against ≤ 0.025 for every aromatic system in the dataset (Table S4, Figure S3). The aromatic LUMOs are ring π^*^ states with nowhere to go; the cysteine LUMO is a backbone state that spreads out exactly as a delocalising orbital should.

For completeness we report the two-term decomposition for cysteine, but it should be read with more caution than the aromatic cases, because the threshold criterion then selects *different kinds of state* on different residues as n grows. In (C)_10_ the per-residue local gaps fall into two families, about 4.7 to 5.2 eV on residues 1, 2, 8 and 10 (backbone π-like) and about 8.1 eV on residues 3, 4, 7 and 9 (σ/lone-pair), with two residues having no state above threshold at all (Table S3). ⟨*g*_*f*_⟩ therefore mixes state characters and its upward drift should not be read as a physical widening of any single excitation energy. The aromatic systems suffer no such ambiguity: there the frontier pair is unambiguously the same ring π/π^*^ pair on every residue, which is why their local gaps form the tight distributions of Figure 5 (for (W)_10_, 3.517 to 3.724 eV across all ten residues).

**Figure 5.**
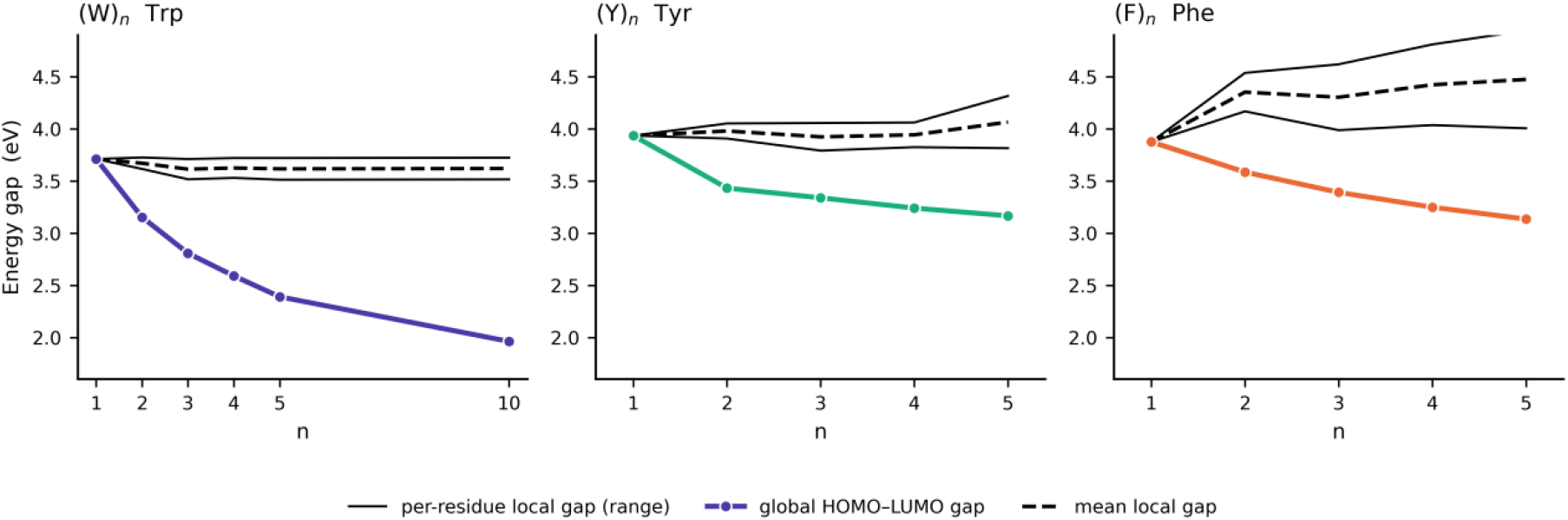
Global HOMO–LUMO gap (coloured line) against the distribution of per-residue local gaps (solid lines, range; dashed line, mean) for Trp, Tyr and Phe. The local gap is flat in n. The global gap is not. The entire chain-length dependence lives in the distance between the two.

**Figure 6.**
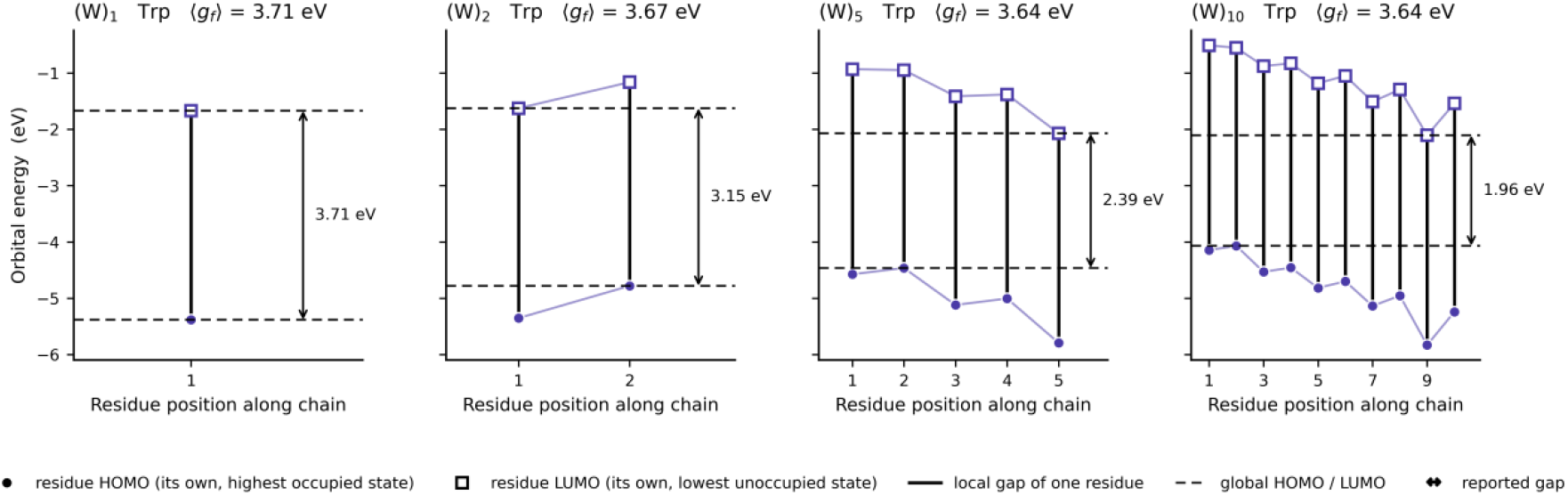
The mechanism for (W)_1_, (W)_2_, (W)_5_ and (W)_10_. Filled circles are each residue’s own HOMO, open squares its own LUMO, and the vertical black segments the local gaps, all the same length in every panel. The site energies slide steadily downward along the chain, so the outermost two (heavy dashed lines) move apart as n grows, and the gap between them, the reported gap, narrows from 3.71 to 1.96 eV.

**Figure 7.**
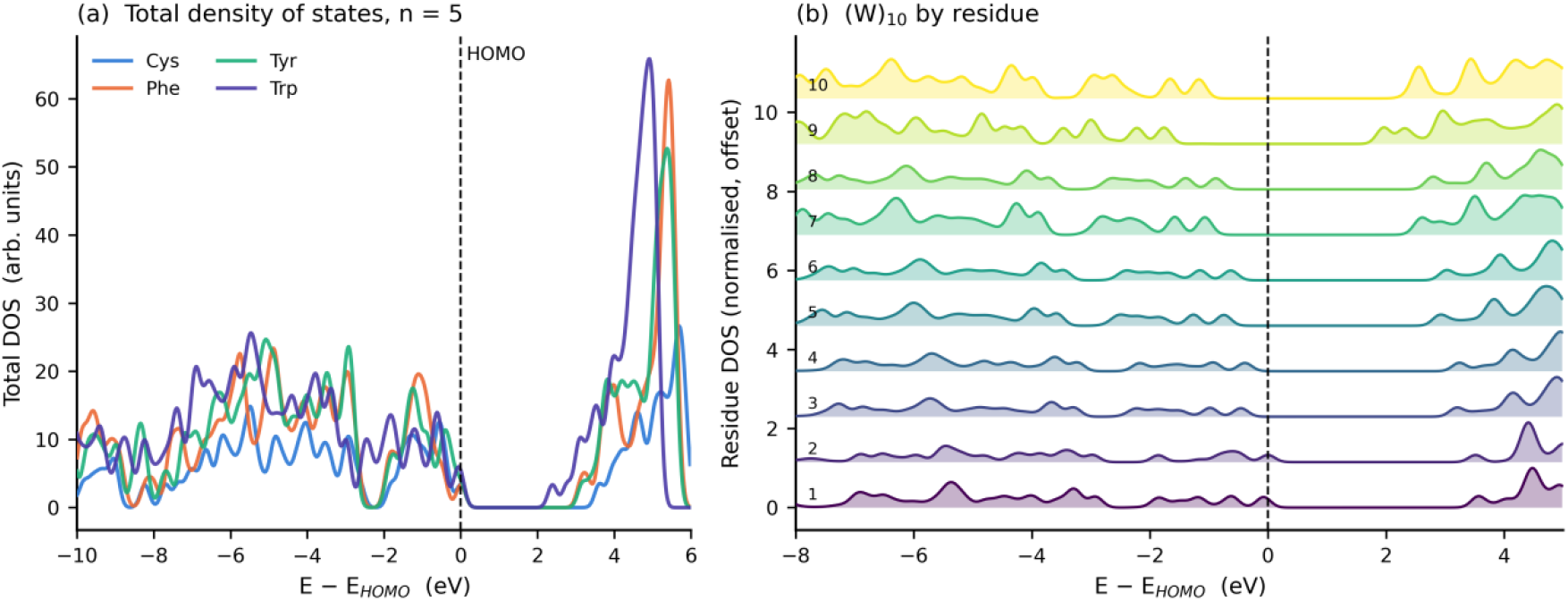
(a) Total density of states for the four n = 5 peptides, each aligned on its own HOMO. (b) The same quantity for (W)_10_, projected onto each residue separately, normalised and offset vertically. The shape of the local density of states is essentially identical for all ten residues; what changes is its position, sliding steadily downhill along the chain. Residue 1 (bottom) keeps the highest occupied states and residue 10 (top) acquires the lowest empty ones: the electrostatic ramp of Section 2.5 seen directly in the electronic structure.

This contrast is the cleanest confirmation that the aromatic behaviour is specifically a *localised-chromophore* effect. When the frontier orbital is a compact ring state that cannot delocalise, chain length can only move it up and down electrostatically, and the extreme-value mechanism operates cleanly and produces a strong linear trend. When the frontier orbital is a backbone state that can delocalise, the two terms of equation (1) grow together and largely cancel, and no trend appears.

One caveat: cysteine’s n = 1 point is anomalous, sitting 0.78 eV below n = 2, because in the single-residue peptide the HOMO is a high-lying thiol sulfur lone pair (the highest HOMO in the dataset) whereas from n = 2 on it is a backbone nitrogen lone pair. The series is not homogeneous across its full range; restricting to n = 2 to 10 gives −0.052 ± 0.014 eV per residue (R = −0.906), a factor of three below the aromatic value and small because the remaining dependence is a backbone effect acting on a delocalised orbital.

### 2.7 What the computed gap measures, and what should be measured instead

The analysis has a consequence that cuts against taking the headline numbers at face value. The global HOMO sits on residue 1 or 2 and the global LUMO on residue n or n−1, separated by up to 32 Å, so the quantity we have been calling “the HOMO–LUMO gap” is the energy difference between two orbitals that barely overlap in space. That is the defining signature of a **charge-transfer-like gap**: the transition would move an electron from one end of the peptide to the other.

Such gaps are the canonical failure case for semilocal DFT. A semilocal functional lacks the derivative discontinuity of the exact one [8, 9], which manifests as delocalisation error: approximate functionals systematically underestimate the energy of spatially separated charge-transfer excitations, often severely [10–14]. In the present case this error acts specifically on Δ(n), the term carrying the entire chain-length dependence, and it grows with the separation of the two edges, so it will inflate the apparent slope. The computed slope of −0.17 to −0.18 eV per residue should therefore be treated as an **upper bound**, not a converged prediction.

More usefully, the analysis implies a testable prediction about the observable experiments actually measure. The lowest *optical* excitation is not the inter-site transition described by the global HOMO–LUMO pair: those orbitals barely overlap, so their direct transition carries almost no dipole strength. It is a **Frenkel exciton** [15, 16] confined to a single aromatic side chain, the localised excitation responsible for the fluorescence of these residues [17], and we have measured directly that each residue’s local gap is invariant with chain length (Section 2.3, Table 2). It follows that:

#### Prediction

The lowest optical excitation energy of NH_2_–(X)_n_–COOH, and hence the position of the absorption edge, should be **nearly independent of n** for X = Phe, Tyr and Trp, even though the Kohn–Sham gap collapses by 1.75 eV from n = 1 to n = 10. The chain-length dependence reported here is a property of the spatially indirect Kohn–Sham gap and should not be read as a redshift of the absorption edge.

This is directly testable by UV absorption or excitation spectroscopy on size-selected oligopeptides, and it is the most valuable experimental consequence of the present work. It also makes an obvious target for a follow-up calculation with a range-separated hybrid functional, in which the inter-site gap is corrected toward its physical value while the local gap is much less affected, precisely the behaviour expected if our interpretation is correct.

### 2.8 Design rules for peptide-based bio-organic semiconductors

Everything above was computed for isolated peptides in vacuum, but the results bear directly on the question that arises whenever a peptide is proposed as an electronic material: what does the backbone itself contribute to charge and energy transport, and what can be designed with it? Five consequences follow, ordered from the most cautionary to the most constructive.

#### A length-dependent gap is not, by itself, evidence of delocalization

Peptide and protein nanostructures, e.g., amyloid fibrils, peptide nanotubes, protein nanowires, are repeatedly reported to be semiconducting [18, 19], and a gap that shifts with chain length or aggregation state is naturally read as the signature of states delocalised along the backbone. Our results show that inference does not follow. Nothing here delocalises, yet the global gap still collapses by 1.19 eV between n = 2 and n = 10 because the two localised edges slide apart on the backbone’s own electrostatic ramp. A length-dependent gap can be produced *entirely* by the electrostatic alignment of localised sites. Because a length-dependent gap and a genuine band edge are not distinguishable from the gap alone, the local-versus-global comparison used here should be a routine control before any length dependence is attributed to delocalisation; it costs one LORBIT = 11 run, far less than the transport measurement it would qualify, and settles the question directly, since a band that forms would show a rising participation number and a growing off-residue weight, neither of which we observe.

#### The site-energy landscape is intrinsic, directional and large

In the standard picture of charge transport in disordered organic semiconductors the site energies are drawn from a Gaussian of width σ ≈ 50 to 150 meV, and that disorder is extrinsic: it comes from packing, impurities and conformational variation, and processing is directed at reducing it [20, 21]. Peptides differ in three ways that change how they should be treated. *The spread is far larger*: the site energies slide monotonically along the chain by a total of 0.4 to 1.4 eV depending on the residue, an order of magnitude above σ for a well-ordered organic semiconductor and comparable to the transport gap itself. *It is systematic, not random*: the correlation with residue index reaches |R| = 0.90 to 1.00, so this is a directed ramp rather than statistical disorder, and the two have different consequences: Gaussian disorder localises carriers in the tails of a distribution, whereas a monotonic ramp biases them in one direction. *It is intrinsic*: the ramp is present in a single isolated molecule in vacuum because it is generated by the backbone’s own permanent dipole field, and no purification, annealing or crystal engineering can remove it. This is what most sharply separates a peptide from a conventional organic semiconductor: the latter’s energetic landscape is a property of how the molecules were assembled, whereas the peptide’s is a property of the molecule itself. The practical consequence is that a charge moving along a peptide always moves downhill or uphill and never on the flat, and the two ends of a chain are electronically inequivalent even when chemically identical.

#### Optical and transport gaps become separately tunable

In a conventional organic semiconductor both gaps are set by the same π system and differ mainly by the exciton binding energy, which is why they can rarely be optimised independently. In a peptide they are set by different parts of the molecule: the **optical** gap is a property of the *side chain*, since the local gap is invariant with chain length to within 0.03 eV for Trp across n = 2 to 10, so the absorption edge is fixed by which aromatic residue is present and little else; the **alignment** gap is a property of the *backbone*, being the extreme-value functional of the shifted site energies, and it responds to chain length, to the charge state of the termini, and to anything else that modifies the internal field. The two knobs are therefore nearly independent, which is not true of a one-chromophore conjugated polymer. A peptide-based material could in principle be tuned for its absorption edge by choosing the residue and for its level alignment, and hence its injection barriers, by choosing the length and charge state of the backbone.

#### The gap-defining state is a dark charge-transfer state with a small binding energy

Since the global HOMO and LUMO sit on different residues, the state whose energy the global gap measures is a charge-transfer state of the kind familiar from organic photovoltaics [21–23]: an electron on one residue and a hole on another. Two properties follow from our numbers. It is **optically dark: its two orbitals** barely overlap, their centroids reaching 32 Å apart by n = 10, so the transition dipole between them is close to zero, and absorption goes instead to a localised Frenkel-like excitation on a single side chain. And its **binding energy is small**: with the edges separated by r ≤ 32 Å and protein-like screening ε_*r*_ ≈ 3 to 4, the Coulomb binding is e^2^/(4πε0ε_*r*_ r) ≈ 0.1 to 0.15 eV, several times below the 0.3 to 0.5 eV typical of molecular charge-transfer states in organic photovoltaic blends [23], because the peptide holds its charges unusually far apart. The peptide is thus a system with poor exciton-to-CT coupling but a low barrier to charge separation once the CT state is reached. That combination, a dark entrance and an easily opened exit, is the specific design opportunity this analysis identifies.

#### Where the backbone does enter the frontier

**cysteine read as a lead**Exactly one system in our set has a backbone contribution to a frontier orbital: the non-aromatic control, cysteine, whose LUMO carries up to 0.38 of its weight off the host residue and whose site HOMO is a sulfur or backbone lone-pair state rather than a ring π orbital. If intrinsic backbone conduction is the goal, this is where to look, because a frontier orbital not confined to a pendant ring is the precondition for a delocalised carrier state, and in our data only the sulfur-containing non-aromatic residue supplies one. This is consistent with the long-standing emphasis on sulfur in protein electron transfer [3, 6] and with the observation that conduction through proteins is generally better than through saturated molecules but worse than through conjugated ones, with cofactors and non-aromatic pathways doing much of the work [24]. It suggests a concrete prediction: cysteine- and methionine-rich sequences, rather than phenylalanine-rich ones, are where a peptide conductor should be sought.

Three caveats bound the reach of all this. The calculations are for single isolated molecules in vacuum; in a fibril or film the neighbouring molecules, the hydration shell and any counter-ions each contribute their own fields, and the internal ramp would be screened and superposed on whatever the packing imposes. Since the total potential drop is roughly constant in n, the per-residue field already dilutes as 1/n, and an external field strong enough to flatten the ramp would remove the chain-length dependence altogether. Solvent in particular should screen the ramp and flatten the trend, so the gas-phase slopes are upper bounds. The gaps are Kohn–Sham gaps from a semilocal functional, which overestimate the chain-length dependence of a spatially indirect gap specifically, and they are not optical gaps. None of this touches the structural result on which the design arguments rest, namely that the frontier orbitals are single-residue objects and that the local gap is invariant with chain length. Those two facts are robust.

## 3. Conclusions

We have computed the Kohn–Sham frontier-orbital gaps of 22 isolated neutral peptides NH_2_–(X)_n_–COOH for X = Phe, Tyr, Trp and Cys, and resolved each state onto individual residues using LORBIT = 11 projections.

The gap narrows with chain length for the three aromatic residues at −0.17 to −0.18 eV per residue, and the raw numbers invite a conjugation-based reading. That reading is wrong. Every aromatic frontier orbital is 97 to 100 % localised on a single residue regardless of chain length, the participation number saturates at one ring, and the mean intra-residue gap is independent of chain length to within 0.03 eV for Trp while the global gap falls by 1.75 eV from n = 1 to n = 10.

The correct mechanism is electrostatic and statistical. The polar backbone generates a monotonic internal potential that shifts each residue’s orbital energies by an amount depending on its position; the global gap is the extreme-value functional min_*f*_ *L*_*f*_ − max_*f*_ *H*_*f*_ of those shifted site energies; and its alignment penalty Δ(n) grows with n because a longer chain places its outermost residues further apart in that potential. The global HOMO and LUMO consequently sit on different residues, up to 32 Å apart, so the narrowing is entirely an inter-site effect with no contribution from any change in the residues themselves. That the slope is independent of chromophore identity follows directly, and the non-aromatic control behaves completely differently: cysteine’s LUMO is delocalised over the backbone (off-residue weight up to 0.38, against ≤ 0.025 for every aromatic system), its HOMO is a sulfur or backbone lone-pair state, and over its full range n = 1 to 10 it shows no significant chain-length trend. Restricting to n = 2 to 10 leaves a small but significant residual slope of −0.052 ± 0.014 eV per residue, a factor of three below the aromatic value and itself a backbone effect acting on a delocalised orbital (Section 2.6).

Three consequences reach beyond this dataset. The first is methodological: for any system in which the frontier orbitals are localised on separate sites, a global Kohn–Sham gap is an inter-site quantity and must be decomposed before it is interpreted; the local-versus-global comparison used here is a cheap and decisive diagnostic that should be applied before any length dependence is attributed to delocalisation. The second is experimental: the chain-length dependence reported here should **not** appear in the absorption spectrum, because the lowest optical excitation is a localised Frenkel exciton on a single side chain whose energy does not depend on chain length. The third is constructive: for bio-organic semiconductors the analysis separates two knobs that are ordinarily entangled, i.e., the side chain sets the absorption edge while the backbone sets the level alignment, and it identifies the backbone’s electrostatic ramp as a directed internal field that is a property of the molecule and therefore cannot be processed away. Realising this programme also requires choosing which sequences to make, a search problem that machine learning has begun to solve: deep learning models now predict the self assembly of short peptides across sequence spaces of more than 10 trillion members [25, 26], human in the loop frameworks have turned such predictions into discovered peptide hydrogels [27], and simple aggregation rules distilled from molecular dynamics simulations now transfer across sequences [28].

## 4. Computational methods

All calculations used VASP 6.1.0 (vasp.6.1.0 28Jan20) [29, 30] with the PBE functional [31] and the projector-augmented-wave method [32, 33], using the standard PAW_PBE potentials N 08Apr2002, C 08Apr2002, O 08Apr2002, H 15Jun2001 and S 06Sep2000.

Each peptide was taken through three sequential stages, all at the Γ point (1 × 1 × 1 Monkhorst-Pack):

1. **bondrelax, geometry optimisation**. A full ionic relaxation with all atoms free to move: conjugate-gradient algorithm (IBRION = 2), force criterion EDIFFG = −1 × 10^−2^ eV Å^−1^, up to NSW = 5000 ionic steps, plane-wave cutoff ENCUT = 600 eV, and EDIFF = 1 × 10^−6^ eV. Gaussian smearing (ISMEAR = 0), NCORE = 4, LREAL = Auto, with VASP’s default symmetry handling retained. Every relaxation reached the required force accuracy.
2. **elecrelax, electronic relaxation**. A single-point SCF calculation at the relaxed geometry (IBRION = −1, NSW = 0) at the same ENCUT = 600 eV but with PREC = Accurate, ADDGRID = .TRUE., EDIFF = 1 × 10^−6^ eV, symmetry switched off (ISYM = 0) so the density is not constrained by the molecular point group, and Gaussian smearing of deliberately narrow width SIGMA = 0.05 eV. This stage writes the CHGCAR and WAVECAR consumed by stage three.
3. **homolumogap, final single point and orbital projection**. A single point restarting from those converged files (ISTART = 1, ICHARG = 0), with a deliberately minimal INCAR: ISMEAR = 0, LORBIT = 11 (site- and lm-projected PROCAR), NCORE = 8, LREAL = Auto. Because ENCUT and EDIFF are not set, VASP applies its defaults: the cutoff falls back to the largest ENMAX in the POTCAR, 400.0 eV for this potential set, and EDIFF to 1 × 10^−4^ eV. The electronic loop is therefore *less* tightly converged than in stage two, but it begins from an already-converged density and converges in 4 to 6 iterations; the final energy change lay between 2.6 × 10^−6^ eV (for (W)_10_) and 7.3 × 10^−5^ eV (for (W)_3_) across the 22 systems, all satisfying the 1 × 10^−4^ eV criterion (Table S1). The eigenvalues analysed here are consequently converged well below the 1 meV precision at which they are quoted.

Separating the work this way lets the expensive relaxation and the tight SCF be performed once, with the projection stage reading an already-converged density.

The calculations were non-spin-polarised (ISPIN = 1), consistent with closed-shell, even-electron neutral peptides. No dipole correction was applied (LDIPOL = .FALSE., IDIPOL = 0). Band eigenvalues and occupations were read from EIGENVAL, and the occupied/empty partition (occupation > 0.5) was cross-checked against E-fermi in the corresponding OUTCAR.

### Systems and cells

The systems are the neutral free peptides NH_2_-(X)_n_-COOH with X ∈ {Phe, Tyr, Trp, Cys} and n = 1 to 5 for each residue, extended to n = 10 for Cys and Trp, giving 22 systems in total, with formulae 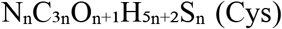, 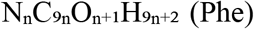, 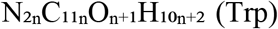 and 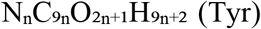, as confirmed from ions per type in the OUTCAR files. Each molecule was placed in an orthorhombic cell with vacuum on all sides, aligned to the Cartesian axes with dimensions growing with the chain so that the vacuum separation is maintained as the molecule lengthens; cell edges range over 14 to 56 Å, 11 to 18 Å and 12 to 51 Å (for example (F)_5_ in 56 × 16 × 51 Å and (W)_10_ in 50 × 15 × 45 Å). Geometries were taken as a single fixed conformer per system, with no conformational sampling, no dispersion correction and no solvent; the consequences are discussed in Section 2.7. The number of valence electrons and bands ranged from 42 in 32 bands for (C)_1_ to 708 in 480 bands for (W)_10_, and the occupied band count recovered from each EIGENVAL file was verified against the electron count implied by the composition and PAW valence charges for all 22 systems, with all 22 agreeing exactly.

### Orbital localisation analysis

Atom- and orbital-resolved projections were obtained from the LORBIT = 11 PROCAR, which gives for every band and ion an lm-decomposed decomposition into s, *p*_*y*_, *p*_*z*_, *p*_*x*_, *d*_*xy*_, *d*_*yz*_, 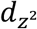, *d*_*xz*_, 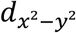 and a total; we use the per-ion total column throughout. To convert per-atom data into chemically meaningful units, atoms were grouped into **residues** by a purely geometric procedure with no reference to the input topology. A heavy-atom bond graph was built by distance criteria with element-pair cutoffs (1.75 Å for C–N, 1.50 Å for C–O, 1.80 Å for C–C, 2.00 Å for C–S, 1.70 Å for N–N), with 5- and 6-membered cycles in the C/N subgraph identifying aromatic rings. Amide C–N bonds, i.e., a C–N contact whose carbon also bears an oxygen within 1.32 Å, were then cut, and the resulting components taken as residues, with hydrogens attached to their nearest heavy atom within 1.55 Å. This recovered exactly n residues for every system with n ≥ 2.

Three localisation metrics were computed from the residue-projected weights: the **participation number** *N*_part_ = (Σ_*i*_ *w*_*i*_)^2^ / Σ_*i*_ *w*_*i*_^2^, the effective number of atoms carrying the orbital, computed on per-atom weights; the **radius of gyration** *R*_*g*_ of the orbital density about its own centroid; and the **HOMO–LUMO centroid separation** d(H–L), the distance between the weight-weighted centroids of the HOMO and LUMO densities. Finally, for each residue f we define its own frontier pair: *H*_*f*_ is the highest occupied band carrying more than a threshold weight fraction on residue f, and *L*_*f*_ the lowest unoccupied band doing so, giving a local gap *g*_*f*_ = *L*_*f*_ − *H*_*f*_. We use a threshold of 0.45 throughout; sensitivity to this choice is discussed in Section 2.3. By construction ⟨*g*_*f*_⟩ ≥ gap_global_, with equality only if a single residue hosts both global edges.

## Data availability

The complete dataset, i.e., eigenvalues, occupations, per-atom lm-decomposed orbital projections, structures and the residue fragmentation, together with all analysis and plotting scripts, is available as peptide_export.json, peptide_allbands.json, peptide_master.csv and the accompanying Python scripts (Tables S7 and S8). Raw VASP input and output files (INCAR, KPOINTS, POSCAR, OUTCAR, EIGENVAL, PROCAR) are archived

## Supplementary information

Tables S1 to S9 and Figures S1 to S3 accompany this paper. They contain the SCF convergence data (Table S1), the threshold-sensitivity scan (Tables S2a and S2b), the per-residue local gaps and site energies (Tables S3 and S5), the off-residue orbital weights (Table S4), the host residues and their separations (Table S6), the complete numerical dataset for all 22 systems (Table S8), the index of figures (Table S9), the reproduction scripts (Table S7), the gap versus HOMO-LUMO separation correlation (Figure S1), the HOMO and LUMO energies separately (Figure S2), and the residue-resolved frontier density of states (Figure S3).

