## Supplementary Information for "Peptide frontier-orbital gaps narrow by backbone electrostatics"

Xu Wu, Wei Fu\*

Shenzhen UniBindMD Computation Biotech Ltd. Co., Shenzhen, Guangdong, P. R. China

##### S1. Geometry optimisation and self-consistency

Each peptide was optimised, then converged, then projected, in three VASP 6.1.0 stages, i.e., bondrelax, elecrelax and homolumogap, as described in Section 4 of the main text.

**Geometry optimisation (bondrelax).** The ionic relaxation ran with IBRION = 2 (conjugate gradient), EDIFFG = $-1 \times 10^{-2}$ eV Å<sup>-1</sup>, NSW = 5000, ENCUT = 600 eV and EDIFF = $1 \times 10^{-6}$ eV. All 22 relaxations terminated with

reached required accuracy - stopping structural energy minimisation

**Electronic relaxation (elecrelax).** The relaxed geometry was then converged with ISTART = 1, ICHARG = 0, ENCUT = 600 eV, PREC = Accurate, ADDGRID = .TRUE., EDIFF = $1 \times 10^{-6}$ eV, ISYM = 0, ISMEAR = 0 and SIGMA = 0.05 eV, writing the CHGCAR and WAVECAR files that the final stage restarts from.

**Final single point (homolumogap).** This stage restarts from those files (ISTART = 1, ICHARG = 0), adds LORBIT = 11 to write the projected PROCAR, and otherwise takes VASP's defaults: ENCUT falls back to the largest ENMAX in the POTCAR (400 eV) and EDIFF to $1 \times 10^{-4}$ eV. The SCF converged in 4 to 6 iterations, and the energy change of the final iteration lay between $2.6 \times 10^{-6}$ and $7.3 \times 10^{-5}$ eV over the 22 systems, every one satisfying the $1 \times 10^{-4}$ eV criterion, and all far below the 1 meV precision at which the eigenvalues are quoted.

**Table S1. The final self-consistency stage, and electron counts**

| System | Atoms | NELECT | NBANDS | Final $\Delta E$ (eV) | SCF iterations |
| --- | --- | --- | --- | --- | --- |
| (C)<sub>1</sub> | 14 | 42 | 32 | $-9.5 \times 10^{-6}$ | 4 |
| (C)<sub>2</sub> | 25 | 76 | 56 | $-2.2 \times 10^{-5}$ | 4 |
| (C)<sub>3</sub> | 36 | 110 | 80 | $-3.3 \times 10^{-5}$ | 4 |
| (C)<sub>4</sub> | 47 | 144 | 96 | $-4.8 \times 10^{-5}$ | 4 |
| (C)<sub>5</sub> | 58 | 178 | 120 | $-6.9 \times 10^{-5}$ | 4 |
| (C)<sub>10</sub> | 113 | 348 | 232 | $-3.0 \times 10^{-5}$ | 5 |
| (F)<sub>1</sub> | 23 | 64 | 48 | $-9.9 \times 10^{-6}$ | 4 |
| (F)<sub>2</sub> | 43 | 120 | 88 | $-2.2 \times 10^{-5}$ | 4 |
| (F)<sub>3</sub> | 63 | 176 | 120 | $-3.7 \times 10^{-5}$ | 4 |
| (F)<sub>4</sub> | 83 | 232 | 160 | $-5.1 \times 10^{-5}$ | 4 |
| (F)<sub>5</sub> | 103 | 288 | 200 | $-6.9 \times 10^{-5}$ | 4 |
| (Y)<sub>1</sub> | 24 | 70 | 48 | $-7.8 \times 10^{-6}$ | 4 |
| (Y)<sub>2</sub> | 45 | 132 | 96 | $-2.1 \times 10^{-5}$ | 4 |
| (Y)<sub>3</sub> | 66 | 194 | 136 | $-4.0 \times 10^{-5}$ | 4 |
| (Y)<sub>4</sub> | 87 | 256 | 176 | $-5.2 \times 10^{-5}$ | 4 |
| (Y)<sub>5</sub> | 108 | 318 | 216 | $-7.1 \times 10^{-5}$ | 4 |
| (W)<sub>1</sub> | 27 | 78 | 56 | $-2.8 \times 10^{-5}$ | 4 |
| (W)<sub>2</sub> | 51 | 148 | 104 | $-4.3 \times 10^{-5}$ | 4 |
| (W)<sub>3</sub> | 75 | 218 | 152 | $-7.3 \times 10^{-5}$ | 4 |
| (W)<sub>4</sub> | 99 | 288 | 200 | $-2.4 \times 10^{-5}$ | 5 |
| (W)<sub>5</sub> | 123 | 358 | 248 | $-3.4 \times 10^{-5}$ | 5 |
| (W)<sub>10</sub> | 243 | 708 | 480 | $-2.6 \times 10^{-6}$ | 6 |

NELECT is the number of valence electrons as reported by VASP, and was independently verified against the composition implied by ions per type and the PAW valence charges (N 5, C 4, O 6, H 1, S 6) for all 22 systems. The number of occupied bands recovered from each EIGENVAL file agreed exactly with NELECT/2 in every case. The final $\Delta E$ column is the energy change of the last self-consistent iteration, taken from the total energy-change (2. order) line of the OUTCAR.

### S2. Sensitivity of the local-gap result to the localisation threshold

The local-gap analysis assigns each residue its own frontier pair $H_f, L_f$ using a threshold THRESH on the residue-projected weight fraction. We scanned THRESH from 0.15 to 0.60 to confirm that the central conclusion, i.e., invariant $\langle g_f \rangle$ and collapsing global gap, is not an artefact of that choice.

**Table S2a. Trp, $\langle g_f \rangle$ drift from n = 2 to n = 10 versus global gap drift**

| THRESH | $\langle g_f \rangle$ at n =<br>2 (eV) | $\langle g_f \rangle$ at n =<br>10 (eV) | local drift<br>(eV) | global drift<br>(eV) | residues used (n =<br>2 / n = 10) |
| --- | --- | --- | --- | --- | --- |
| 0.15 | 3.624 | 3.643 | +0.019 | -1.192 | 2 / 10 |
| 0.20 | 3.624 | 3.643 | +0.019 | -1.192 | 2 / 10 |
| 0.25 | 3.671 | 3.643 | -0.028 | -1.192 | 2 / 10 |
| 0.30 | 3.671 | 3.643 | -0.028 | -1.192 | 2 / 10 |
| 0.35 | 3.671 | 3.643 | -0.028 | -1.192 | 2 / 10 |
| 0.40 | 3.671 | 3.643 | -0.028 | -1.192 | 2 / 10 |
| **0.45** | **3.671** | **3.643** | **-0.028** | **-1.192** | **2 / 10** |
| 0.50 | 3.671 | 3.742 | +0.071 | -1.192 | 2 / 10 |
| 0.55 | 3.671 | 3.825 | +0.154 | -1.192 | 2 / 10 |
| 0.60 | 3.671 | 3.744 | +0.073 | -1.192 | 2 / 9 |

Over the whole scanned range the local-gap drift stays between $-0.028$ and $+0.154$ eV while the global gap falls by $1.192$ eV, i.e., between 2 % and 13 % of the effect. At every threshold up to 0.55 all ten residues of $(W)_{10}$ host their own well-defined frontier pair, so the analysis discards nothing.

**Table S2b. Local drift as a fraction of global drift, all residues**

Values are (local-gap drift) / (global-gap drift) × 100 %, comparing the longest chain to $n = 2$ for each residue. A value near 0 % means the local gap does not move and the conclusion holds.

| THRESH | Cys | Phe | Tyr | Trp |
| --- | --- | --- | --- | --- |
| 0.15 | -2 % | -29 % | -6 % | -2 % |
| 0.20 | +17 % | -6 % | -6 % | -2 % |
| 0.25 | -70 % | -6 % | -6 % | +2 % |
| 0.30 | -88 % | -6 % | -6 % | +2 % |
| 0.35 | -283 % | -8 % | -6 % | +2 % |
| 0.40 | -286 % | -30 % | -6 % | +2 % |
| **0.45** | -376 % | -44 % | -29 % | **+2 %** |
| 0.50 | -465 % | -71 % | -29 % | -6 % |
| 0.55 | -207 % | -71 % | -29 % | -13 % |
| 0.60 | -88 % | -122 % | -29 % | -6 % |

For Trp the ratio is negligible everywhere in the scanned range ($\leq 13$ % in magnitude, and +2 % at the working threshold). For Tyr it reaches 29 % and for Phe 44 % at the working threshold, and larger at coarser ones; Phe is the extreme case because its local gap genuinely widens with $n$, the opposite sign to the global trend. In no case does the local drift carry the global trend. Cysteine is the exception and is discussed in Section 2.6 of the main text: because its frontier states are not a single well-defined chromophore pair, the threshold picks up different state characters on different residues, and the ratio becomes ill-conditioned once $\langle g_f \rangle$ itself drifts. This is exactly why cysteine is used as a qualitative control on the *presence* of a trend rather than as a quantitative test of equation (1).

#### S3. Per-residue site energies

**Table S3. Each residue's own frontier pair and local gap (THRESH = 0.45)**

| System | Residue | own HOMO (eV) | own LUMO (eV) | local gap (eV) |
| --- | --- | --- | --- | --- |
| (W)<sub>10</sub> | 1 | -4.144 | -0.509 | 3.635 |
| (W)<sub>10</sub> | 2 | -4.068 | -0.552 | 3.517 |
| (W)<sub>10</sub> | 3 | -4.531 | -0.877 | 3.654 |
| (W)<sub>10</sub> | 4 | -4.455 | -0.828 | 3.628 |
| (W)<sub>10</sub> | 5 | -4.816 | -1.181 | 3.635 |
| (W)<sub>10</sub> | 6 | -4.698 | -1.050 | 3.648 |
| (W)<sub>10</sub> | 7 | -5.136 | -1.509 | 3.627 |
| (W)<sub>10</sub> | 8 | -4.952 | -1.294 | 3.659 |
| (W)<sub>10</sub> | 9 | -5.831 | -2.107 | 3.724 |
| (W)<sub>10</sub> | 10 | -5.240 | -1.539 | 3.701 |
| (Y)<sub>5</sub> | 1 | -4.929 | -1.113 | 3.816 |
| (Y)<sub>5</sub> | 2 | -5.142 | -0.825 | 4.316 |
| (Y)<sub>5</sub> | 3 | -5.251 | -1.176 | 4.076 |
| (Y)<sub>5</sub> | 4 | -5.348 | -1.330 | 4.018 |
| (Y)<sub>5</sub> | 5 | -5.822 | -1.762 | 4.060 |
| (F)<sub>5</sub> | 1 | -5.153 | -1.147 | 4.006 |
| (F)<sub>5</sub> | 2 | -5.731 | -1.204 | 4.527 |
| (F)<sub>5</sub> | 3 | -6.363 | -1.419 | 4.944 |
| (F)<sub>5</sub> | 4 | -5.872 | -1.419 | 4.453 |
| (F)<sub>5</sub> | 5 | -6.841 | -2.018 | 4.824 |
| (C)<sub>10</sub> | 1 | -5.226 | -0.522 | 4.704 |
| (C)<sub>10</sub> | 2 | -5.561 | -0.439 | 5.122 |
| (C)<sub>10</sub> | 3 | -8.152 | -0.065 | 8.087 |
| (C)<sub>10</sub> | 4 | -8.210 | -0.116 | 8.095 |
| (C)<sub>10</sub> | 7 | -8.344 | -0.278 | 8.067 |
| (C)<sub>10</sub> | 8 | -6.053 | -0.893 | 5.160 |
| (C)<sub>10</sub> | 9 | -8.446 | -0.293 | 8.153 |
| (C)<sub>10</sub> | 10 | -6.874 | -1.765 | 5.109 |

Two features deserve comment.

For (W)<sub>10</sub> the ten local gaps span only 3.517 to 3.724 eV, a spread of 0.21 eV, while the site HOMO energies span 1.76 eV and the site LUMO energies 1.60 eV. The site energies move by nearly an order of magnitude more than the gaps do, which is the direct signature of a rigid, common-mode electrostatic shift, and is the observation on which the mechanism rests.

For (C)<sub>10</sub> the local gaps fall into two clearly separated families, ~4.7 to 5.2 eV (residues 1, 2, 8, 10) and ~8.1 eV (residues 3, 4, 7, 9), with residues 5 and 6 absent because no state there exceeds the 45 % threshold. This confirms the caveat in Section 2.6 of the main text:

for cysteine the criterion is selecting different *kinds* of state (backbone π-like versus $\sigma$/lone-pair) on different residues. The apparent widening of $\langle g_f \rangle$ with $n$ for cysteine is therefore partly a change in which states are being compared, not a physical widening of a single excitation energy. The aromatic systems show no such ambiguity.

### S4. Delocalisation of the frontier orbitals

**Table S4. Off-residue weight of the global HOMO and LUMO**

The weight fraction of each frontier orbital that lies outside the single residue carrying the largest share. A delocalised orbital approaches 1; a perfectly localised one is 0.

| System | HOMO off-residue | LUMO off-residue |
| --- | --- | --- |
| (C)<sub>1</sub> | 0.3943 | 0.4095 |
| (C)<sub>2</sub> | 0.0138 | 0.1601 |
| (C)<sub>3</sub> | 0.0204 | 0.2826 |
| (C)<sub>4</sub> | 0.0204 | 0.3267 |
| (C)<sub>5</sub> | 0.0221 | 0.3472 |
| (C)<sub>10</sub> | 0.0256 | **0.3772** |
| (F)<sub>1</sub> | 0.0000 | 0.0000 |
| (F)<sub>2</sub> | 0.0116 | 0.0167 |
| (F)<sub>3</sub> | 0.0147 | 0.0229 |
| (F)<sub>4</sub> | 0.0148 | 0.0229 |
| (F)<sub>5</sub> | 0.0180 | 0.0248 |
| (Y)<sub>1</sub> | 0.0000 | 0.0000 |
| (Y)<sub>2</sub> | 0.0069 | 0.0044 |
| (Y)<sub>3</sub> | 0.0134 | 0.0066 |
| (Y)<sub>4</sub> | 0.0101 | 0.0065 |
| (Y)<sub>5</sub> | 0.0101 | 0.0087 |
| (W)<sub>1</sub> | 0.0000 | 0.0000 |
| (W)<sub>2</sub> | 0.0075 | 0.0021 |
| (W)<sub>3</sub> | 0.0110 | 0.0043 |
| (W)<sub>4</sub> | 0.0074 | 0.0043 |
| (W)<sub>5</sub> | 0.0111 | 0.0043 |
| (W)<sub>10</sub> | 0.0203 | 0.0043 |

The aromatic LUMOs carry at most 2.5 % of their weight off-residue at any chain length, and the aromatic HOMOs at most 2.0 %. Cysteine's LUMO reaches 37.7 %. The value for the $n = 1$ system (C)<sub>1</sub> is not physically meaningful: with only one residue the geometric fragmentation necessarily separates the capping NH<sub>2</sub> and COOH groups, which inflates the “off-residue” fraction for the capping-group states. The (F)<sub>1</sub>, (Y)<sub>1</sub> and (W)<sub>1</sub> entries are exactly 0.0000 because in those cases the single residue is the only fragment hosting the frontier orbital.

### S5. The two-term decomposition

**Table S5. Mean intra-residue gap, its spread, and the alignment penalty**

$\Delta(n) = \langle g_f \rangle - \text{gap}_{\text{global}}$ is the alignment penalty of equation (1) of the main text.

| System | n | $\langle g_f \rangle$ (eV) | range of $g_f$ (eV) | global gap (eV) | $\Delta$ (eV) | $\Delta/\langle g_f \rangle$ |
| --- | --- | --- | --- | --- | --- | --- |
| (C)<sub>1</sub> | 1 | 4.306 | 3.775 to 4.837 | 3.156 | 1.150 | 26.7 % |
| (C)<sub>2</sub> | 2 | 4.765 | 4.622 to 4.909 | 3.939 | 0.827 | 17.3 % |
| (C)<sub>3</sub> | 3 | 4.931 | 4.668 to 5.148 | 3.772 | 1.159 | 23.5 % |
| (C)<sub>4</sub> | 4 | 5.129 | 4.699 to 5.840 | 3.657 | 1.472 | 28.7 % |
| (C)<sub>5</sub> | 5 | 5.675 | 4.698 to 7.977 | 3.608 | 2.068 | 36.4 % |
| (C)<sub>10</sub> | 10 | 6.562 | 4.704 to 8.153 | 3.461 | 3.101 | 47.3 % |
| (F)<sub>1</sub> | 1 | 3.876 | 3.876 to 3.876 | 3.876 | 0.000 | 0.0 % |
| (F)<sub>2</sub> | 2 | 4.353 | 4.169 to 4.538 | 3.586 | 0.767 | 17.6 % |
| (F)<sub>3</sub> | 3 | 4.402 | 3.988 to 4.620 | 3.392 | 1.010 | 22.9 % |
| (F)<sub>4</sub> | 4 | 4.473 | 4.037 to 4.811 | 3.249 | 1.223 | 27.4 % |
| (F)<sub>5</sub> | 5 | 4.551 | 4.006 to 4.944 | 3.135 | 1.416 | 31.1 % |
| (Y)<sub>1</sub> | 1 | 3.936 | 3.936 to 3.936 | 3.936 | 0.000 | 0.0 % |
| (Y)<sub>2</sub> | 2 | 3.980 | 3.908 to 4.053 | 3.433 | 0.548 | 13.8 % |
| (Y)<sub>3</sub> | 3 | 3.966 | 3.792 to 4.057 | 3.339 | 0.627 | 15.8 % |
| (Y)<sub>4</sub> | 4 | 3.984 | 3.825 to 4.062 | 3.241 | 0.743 | 18.6 % |
| (Y)<sub>5</sub> | 5 | 4.057 | 3.816 to 4.316 | 3.167 | 0.891 | 22.0 % |
| (W)<sub>1</sub> | 1 | 3.712 | 3.712 to 3.712 | 3.712 | 0.000 | 0.0 % |
| (W)<sub>2</sub> | 2 | 3.671 | 3.616 to 3.726 | 3.153 | 0.518 | 14.1 % |
| (W)<sub>3</sub> | 3 | 3.642 | 3.517 to 3.711 | 2.808 | 0.834 | 22.9 % |
| (W)<sub>4</sub> | 4 | 3.648 | 3.530 to 3.721 | 2.590 | 1.058 | 29.0 % |
| (W)<sub>5</sub> | 5 | 3.641 | 3.513 to 3.721 | 2.390 | 1.251 | 34.4 % |
| (W)<sub>10</sub> | 10 | 3.643 | 3.517 to 3.724 | 1.962 | 1.681 | 46.1 % |

### S6. Where the global HOMO and LUMO sit

**Table S6. Host residues and their separation**

| System | HOMO host<br>residue | weight | LUMO host<br>residue | weight | centroid separation<br>(Å) |
| --- | --- | --- | --- | --- | --- |
| (C)<sub>2</sub> | 1 | 0.99 | 2 | 0.84 | 4.42 |
| (C)<sub>3</sub> | 1 | 0.98 | 3 | 0.72 | 7.13 |
| (C)<sub>4</sub> | 1 | 0.98 | 4 | 0.67 | 10.67 |
| (C)<sub>5</sub> | 1 | 0.98 | 5 | 0.65 | 13.98 |
| (C)<sub>10</sub> | 1 | 0.97 | 10 | 0.62 | 31.67 |
| (F)<sub>2</sub> | 1 | 0.99 | 2 | 0.98 | 6.65 |
| (F)<sub>3</sub> | 1 | 0.99 | 3 | 0.98 | 9.55 |
| (F)<sub>4</sub> | 1 | 0.99 | 3 | 0.98 | 13.12 |
| (F)<sub>5</sub> | 1 | 0.98 | 5 | 0.98 | 16.43 |
| (Y)<sub>2</sub> | 2 | 0.99 | 1 | 1.00 | 7.33 |
| (Y)<sub>3</sub> | 1 | 0.99 | 3 | 0.99 | 8.78 |
| (Y)<sub>4</sub> | 1 | 0.99 | 3 | 0.99 | 12.92 |
| (Y)<sub>5</sub> | 1 | 0.99 | 5 | 0.99 | 15.64 |
| (W)<sub>2</sub> | 2 | 0.99 | 1 | 1.00 | 8.49 |
| (W)<sub>3</sub> | 2 | 0.99 | 3 | 1.00 | 7.64 |
| (W)<sub>4</sub> | 2 | 0.99 | 3 | 1.00 | 13.16 |
| (W)<sub>5</sub> | 2 | 0.99 | 5 | 1.00 | 14.50 |
| (W)<sub>10</sub> | 2 | 0.98 | 9 | 1.00 | 32.71 |

The pattern is systematic in every system with $n \geq 3$: the global HOMO is hosted by the residue at the *low* end of the chain index (residue 1 or 2) and the global LUMO by the residue at the *high* end (residue $n$ or $n-1$). The two $n = 2$ systems, (W)<sub>2</sub> and (Y)<sub>2</sub>, are the exception and are reversed, i.e., their HOMO sits on residue 2 and their LUMO on residue 1, so the ordering of the two edges is not fixed at $n = 2$, only their separation. The separation grows with $n$, reaching $\approx 32$ Å at $n = 10$. The trend is systematic, though not strictly monotonic ((W)<sub>3</sub> sits slightly below (W)<sub>2</sub>). The global HOMO and LUMO are therefore never on the same residue from $n = 2$ onward, which is the structural origin of the alignment penalty $\Delta(n)$.

### S7. Gap versus HOMO–LUMO separation

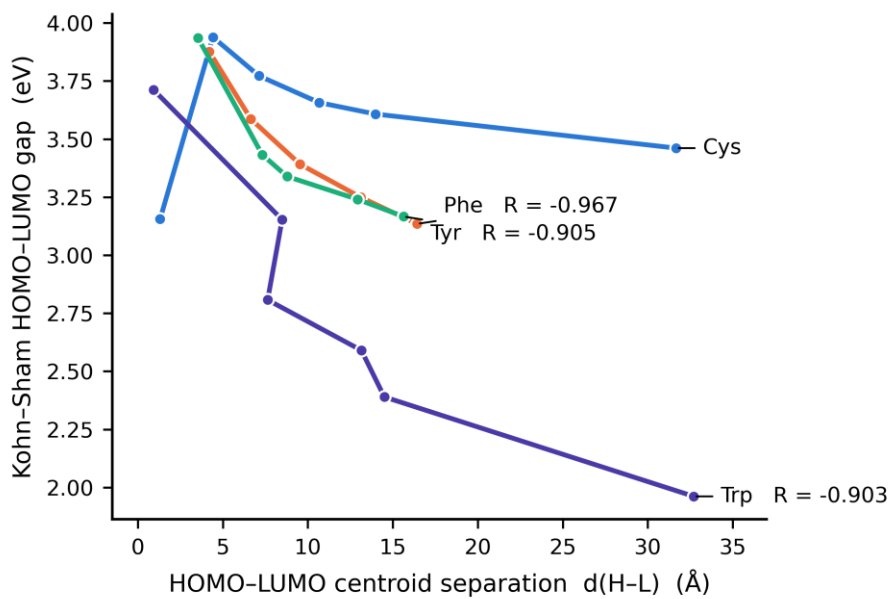

**Figure S1.** Kohn–Sham gap versus the separation of the HOMO and LUMO centroids. The gap falls as the two edges are pulled apart, with Pearson $R = -0.967$ (Phe), $-0.905$ (Tyr) and $-0.903$ (Trp) over all chain lengths, and $R = -0.988$, $-0.984$ and $-0.896$ respectively when the $n = 1$ points are excluded. Cysteine is shown for reference and is much flatter ($R = -0.099$ over all points). The correlation is the signature of a spatially indirect gap: the further apart the two edges are, the lower the energy difference between them.

### S8. The HOMO and LUMO energies separately

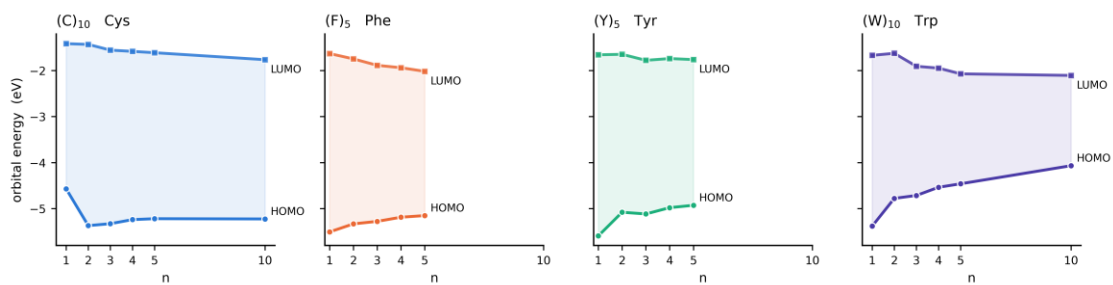

**Figure S2.** The HOMO and LUMO energies separately. Both edges move as $n$ grows, but by different amounts: the HOMO rises (becomes less bound) and the LUMO falls, so their separation changes rather than staying fixed. For Trp the HOMO supplies about three quarters of the total change; for Phe the two contributions are of similar size and opposite sign, so the HOMO dominance is not universal across the three residues and should not be over-read. The reliable content of this observation is one of accounting: the closing of the gap is not carried by one edge alone, so a mechanism that shifts the whole frontier manifold rigidly cannot be the whole story.

### S9. The frontier region of the density of states, resolved by residue

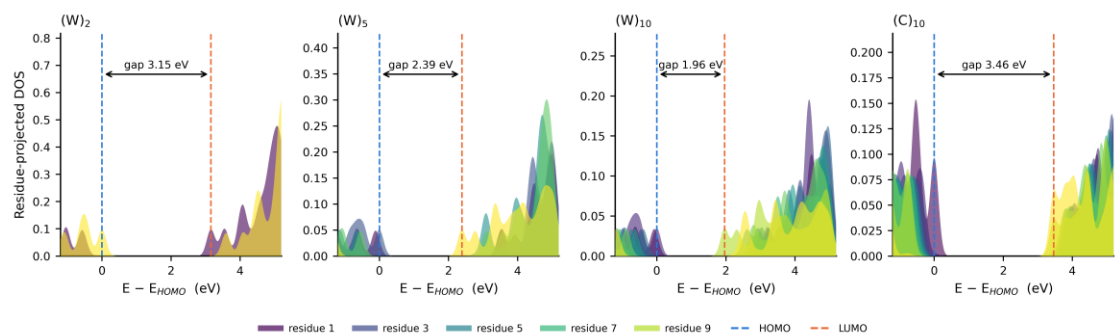

**Figure S3.** The frontier region of the density of states, resolved by residue, for $(W)_2$, $(W)_5$ and $(W)_{10}$, with $(C)_{10}$ shown for contrast. Each panel is normalised by the maximum of its own total DOS so the areas are comparable between panels. For the three Trp peptides a single occupied peak sits just below the dashed blue line and a single empty peak just above the dashed orange line; those two lines are the global HOMO and LUMO. The peak at the LUMO edge is carried by the highest-numbered residues (yellow) and the peak at the HOMO edge by the lowest-numbered ones (dark), so the two edges are drawn from opposite ends of the chain, which is exactly what makes $\Delta(n)$ grow with $n$. Cysteine behaves differently: its empty edge is not a single residue's peak but a broad distribution with substantial weight from residues 7, 8, 9 and 10 at once, the delocalisation that is the reason its gap shows no trend.

### S10. Reproduction

#### Table S7. Reproduction scripts

All analysis scripts and intermediate data are provided alongside this document:

| File | Contents |
| --- | --- |
| extract_gap.py | EIGENVAL/OSZICAR parsing; HOMO, LUMO, gap |
| procar_analysis.py | element- and orbital-resolved frontier-orbital character |
| analyze2.py | residue fragmentation by amide-bond cutting |
| export.py | exports coordinates, elements, energies and a 27-band PROCAR window |
| export2.py | exports all-band per-atom PROCAR totals (basis for the local-gap analysis) |
| localgap.py | local versus global gap |
| mechanism.py | site-energy extraction and the $\Delta(n)$ decomposition |
| final_model.py | ramp fits, $\langle g_f \rangle$ drift, alignment penalty |
| ramp_or_disorder.py | discrimination between a systematic ramp and statistical disorder |
| thresh_scan.py | the threshold scan of Section S2 |
| checks.py | slope robustness, gap versus d(H–L) correlations, master CSV export |
| plots.py, fig6.py, si_fig.py | figures 1, 4, 5, 6, 7 and S1 (matplotlib, 600 dpi) |
| figs_extra.py | figures 7 and S3 (DOS and frontier-region panels) |
| pymol_figs.py | figures 2 and 3: writes PDB files from the JSON, ray-traces one panel per molecule with PyMOL at a common Å-per-pixel scale, and composites the panels at 600 dpi |
| peptide_export.json | coordinates, elements, eigenvalues, PROCAR window (2.5 MB) |
| peptide_allbands.json | all-band per-atom weights, fragmentation, eigenvalues (2.0 MB) |
| peptide_master.csv | the master data table |

Raw VASP files (INCAR, KPOINTS, POSCAR, OUTCAR, EIGENVAL, PROCAR) are archived for all 22 systems under `homolumogap/<Residue>_homolumogap/<n><Residue>/` on the computing facility. Additional `*partialcharge` directories present in the archive contain `LPARD = .TRUE.` partial-charge-density runs (PARCHG) of the *same* neutral structures, generated for band-density visualisation; they are post-processing of the same wavefunctions and are correctly excluded from all statistics reported here, since including them would double-count systems.

### S11. Complete dataset

**Table S8. Kohn–Sham frontier levels, gap, HOMO–LUMO centroid separation and HOMO localisation for all 22 systems.**

| System | N_atoms | HOMO (eV) | LUMO (eV) | Gap (eV) | d(H–L) (Å) | HOMO $N_{\text{part}}$ | $R_g(\text{HOMO})$ (Å) |
| --- | --- | --- | --- | --- | --- | --- | --- |
| (C)<sub>1</sub> | 14 | –4.571 | –1.416 | 3.156 | 1.31 | 3.03 | 1.87 |
| (C)<sub>2</sub> | 25 | –5.371 | –1.432 | 3.939 | 4.42 | 3.33 | 2.03 |
| (C)<sub>3</sub> | 36 | –5.329 | –1.557 | 3.772 | 7.13 | 3.39 | 2.02 |
| (C)<sub>4</sub> | 47 | –5.240 | –1.583 | 3.657 | 10.67 | 3.34 | 2.02 |
| (C)<sub>5</sub> | 58 | –5.220 | –1.613 | 3.608 | 13.98 | 3.36 | 2.03 |
| (C)<sub>10</sub> | 113 | –5.226 | –1.765 | 3.461 | 31.67 | 3.53 | 2.08 |
| (F)<sub>1</sub> | 23 | –5.507 | –1.631 | 3.876 | 4.21 | 1.76 | 1.26 |
| (F)<sub>2</sub> | 43 | –5.333 | –1.747 | 3.586 | 6.65 | 2.52 | 1.87 |
| (F)<sub>3</sub> | 63 | –5.278 | –1.887 | 3.392 | 9.55 | 2.43 | 1.73 |
| (F)<sub>4</sub> | 83 | –5.188 | –1.939 | 3.249 | 13.12 | 2.50 | 1.77 |
| (F)<sub>5</sub> | 103 | –5.153 | –2.018 | 3.135 | 16.43 | 2.45 | 1.76 |
| (Y)<sub>1</sub> | 24 | –5.594 | –1.658 | 3.936 | 3.53 | 4.16 | 2.61 |
| (Y)<sub>2</sub> | 45 | –5.080 | –1.647 | 3.433 | 7.33 | 6.66 | 3.12 |
| (Y)<sub>3</sub> | 66 | −5.117 | −1.778 | 3.339 | 8.78 | 4.40 | 2.93 |
| (Y)<sub>4</sub> | 87 | −4.981 | −1.740 | 3.241 | 12.92 | 4.77 | 2.98 |
| (Y)<sub>5</sub> | 108 | −4.929 | −1.762 | 3.167 | 15.64 | 4.67 | 2.94 |
| (W)<sub>1</sub> | 27 | −5.382 | −1.670 | 3.712 | 0.93 | 7.95 | 2.36 |
| (W)<sub>2</sub> | 51 | −4.779 | −1.626 | 3.153 | 8.49 | 8.70 | 2.67 |
| (W)<sub>3</sub> | 75 | −4.715 | −1.908 | 2.808 | 7.64 | 8.34 | 2.80 |
| (W)<sub>4</sub> | 99 | −4.538 | −1.948 | 2.590 | 13.16 | 8.57 | 2.73 |
| (W)<sub>5</sub> | 123 | −4.461 | −2.071 | 2.390 | 14.50 | 8.58 | 2.78 |
| (W)<sub>10</sub> | 243 | −4.068 | −2.107 | 1.962 | 32.71 | 8.95 | 2.84 |

### S12. Index of figures

**Table S9. Index of figures**

| Figure | File | Content |
| --- | --- | --- |
| 1 | figs/fig1_gap_vs_n.png | Gap versus n for all four residues |
| 2 | figs/fig7_structures.png | The peptide structures, with the HOMO- and LUMO-hosting residues coloured (PyMOL) |
| 3 | figs/fig4_orbitals.png | Frontier-orbital densities along the Trp series (PyMOL) |
| 4 | figs/fig5_localisation.png | HOMO participation number versus n |
| 5 | figs/fig3_local_vs_global.png | Local versus global gap |
| 6 | figs/fig6_mechanism.png | Site energies and the extreme-value gap |
| 7 | figs/fig8_dos.png | Total and residue-projected density of states |
| S1 | figs/figS1_gap_vs_dHL.png | Gap versus HOMO-LUMO centroid separation |
| S2 | figs/fig2_levels.png | HOMO and LUMO energies separately |
| S3 | figs/fig9_frontier.png | Frontier region of the DOS, resolved by residue |

Figure numbers run in order of citation in the main text; the file names are descriptive and do not always match the numbers. Figures 3 and 4 are rendered from the relaxed coordinates with PyMOL and composited at 600 dpi; all figures are 600 dpi.
